# Integration of UAS-based spatial surveys and bio-logging tracking enhances precision in population size estimation

**DOI:** 10.64898/2026.01.25.701645

**Authors:** Sota Inoue, Yuichi Mizutani, Hibiki Sugiyama, Yusuke Goto, Ken Yoda

## Abstract

Accurately estimating wildlife population sizes, essential for ecological theory and conservation management, yet remains challenging. Although unmanned aerial systems (UASs) combined with machine learning, have revolutionized population estimation, they face limitations in addressing the hierarchical population processes from individual behavior to colony-and population-level dynamics. To overcome this limitation, we developed a data integration framework that jointly analyzes multiple datasets, representing different scales of the same underlying process, were jointly analyzed. Using a seabird colony as a model system, we integrated UAS-based count data in the colony with bio-logging-based tracking data to estimate population size by quantifying both the number of individuals present and the proportion absent from the surveyed area. These complementary datasets were linked using state-space models allowing accurate population estimates with explicit uncertainty quantification. Furthermore, we evaluated the robustness of the estimations with respect to sample size. Sub-sampling simulations revealed that estimation uncertainty was more sensitive to sample size in bio-logging-based tracking data than in UAS-based count data. This finding highlights the importance of understanding dataset-specific properties when designing effective investigations. Overall, our resource-efficient framework is broadly applicable across species and populations and demonstrates how integrating complementary observation methods can improve population estimates and inform conservation practice.

## Introduction

Population size or abundance serves as an important indicator for ecological theories related to demography, as well as for conservation planning on a regional scale that considers multiple species and populations (Callaghan et al., 2024; Rockwood, 2015). Moreover, sustainable monitoring over the mid-to long-term is essential in the rapidly changing global environment (Reif, 2013). However, long-term monitoring demands significant effort, and even gaining an understanding of the current situation can be challenging, as wild animals often move extensively and inhabit areas that are inaccessible to humans.

Over the past decade, unmanned aerial systems (UASs) have become standard techniques for counting and estimating the abundance of wild animals (Anderson & Gaston, 2013; Robinson et al., 2022). UAS-based aerial census is capable of extensive coverage across diverse environments, including both terrestrial landscapes and marine seascapes, facilitating efficient data collection (Gallego & Sarasola, 2021; Hodgson et al., 2013). Aerial censuses at high altitudes may minimize disturbances compared with ground-based observations within habitats (Krause et al., 2021). Aerial photography often allows higher detection rates than ground-based observations because it reduces the occurrence of individuals overlapping or obstruction by vegetation (Hodgson et al., 2018; McCarthy et al., 2023; Rush et al., 2018). This technology can address a wide range of environments and taxonomic groups, from terrestrial and marine mammals (Johnston et al., 2017; Sweeney et al., 2016) to relatively small birds (Dunn et al., 2021; Sardà-Palomera et al., 2017) and even animal nests (van Andel et al., 2015).

Deep learning-based object detection facilitates the handling of large datasets collected by UASs. This technique is effective for aerial images, where backgrounds are complex and light conditions are unstable, decreasing human effort and observer bias (Hayes et al., 2021; Weinstein et al., 2022). The joint deployment of UAS-based aerial census and deep learning-based object detection upscale data collection and analysis, thereby greatly advancing our understanding of population dynamics (Corcoran et al., 2021; Tuia et al., 2022).

Integration with other data sources can further advance UAS-based aerial census. It is widely recognized that integrating multiple datasets that capture different scales or aspects of the same underlying process offers deeper insights than treating them independently (Kéry & Royle, 2020; Matthiopoulos et al., 2022). Data integration can allow one dataset to validate parameter estimation from the other dataset (Boback et al., 2020), improve the precision of parameter estimates (Blackwell & Matthiopoulos, 2024), and enable the estimation of parameters that cannot be identified from a single dataset alone. Spatial survey data is often integrated with tracking data to estimate animal movement, population distribution, and species distribution (Blackwell & Matthiopoulos, 2024; Kendall et al., 2023; Lauret et al., 2025; Louzao et al., 2009; Yamamoto et al., 2015). In population estimation, spatial count data is often integrated with presence data based on individual-level movement data to complement the limitation of each method. Spatial censuses allow count a large number of individuals within limited space, whereas movement data enables detailed tracking of individuals but is typically constrained by small sample size (Figure 1).

**Figure 1.**
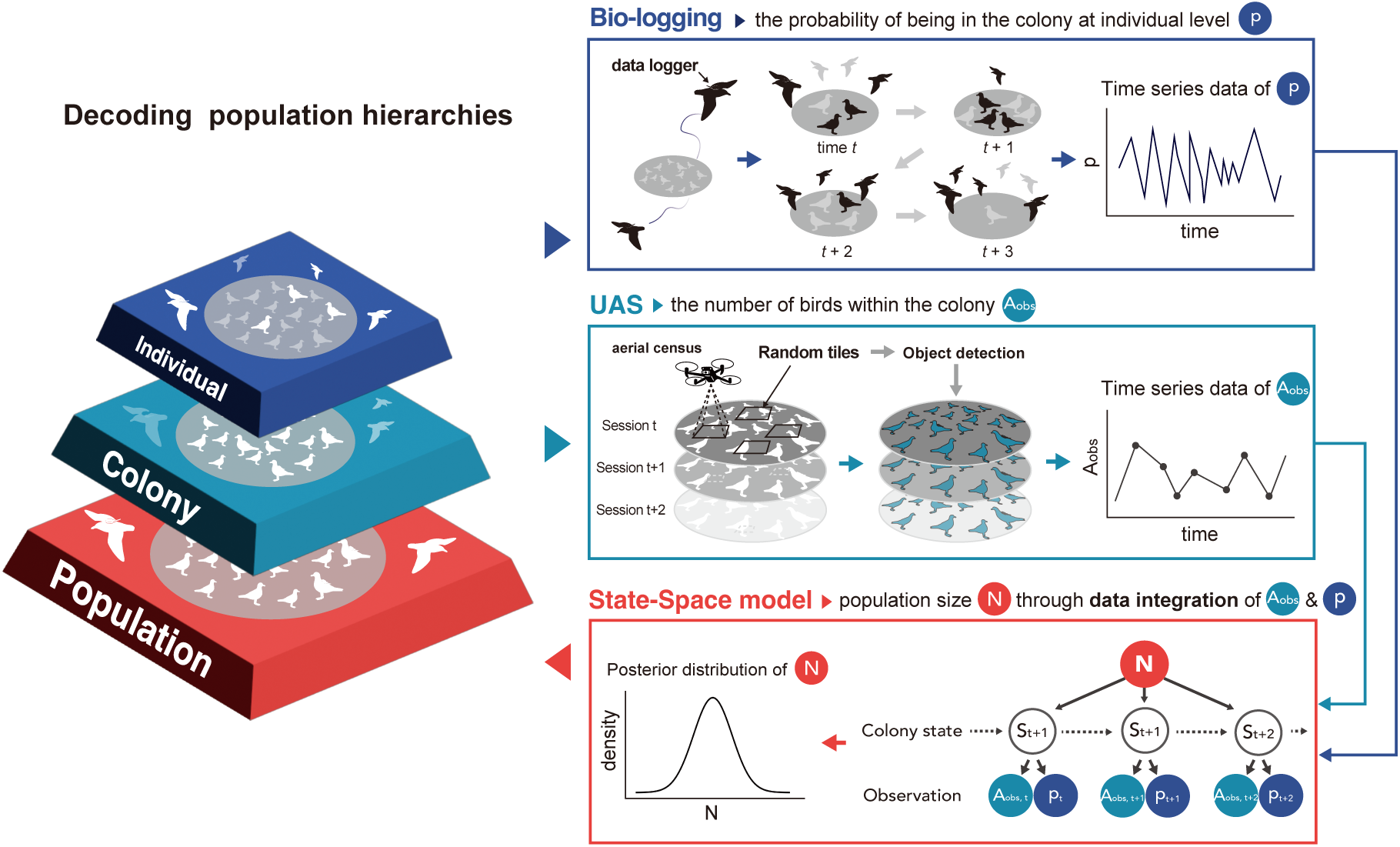
Schematic diagram of the proposed framework, which fuses colony-level census data and individual-level behavior data to estimate the population size *N*. The proportion of individuals, *p*, inside and outside the colony at any given time can be estimated by bio-logging. Individuals present within the colony at any given time are defined as the observable number, *A_obs_*. Unmanned aerial systems (UASs) and deep learning were combined to estimate *A_obs_* at a given time. These time-series data were integrated using a state-space model to estimate the population size.

Cost-effective data collection methods, such as UASs and bio-logging, are likely to become increasingly popular in practical management. Despite their growing popularity, the integration of these data sources has not been systematically investigated. In particular, their potential synergy and methodological challenges have remained largely unexamined.

Accordingly, it will be valuable to explore a generalized framework to integrate these different data sets, as such integration may enable the estimation of absolute population size rather than relative abundance.

Here, we propose a novel framework in which bio-logging-based tracking data complement UAS-based count data by enabling the estimation of individual-level absence. Our aim was to establish an efficient method for estimating population size by integrating data across hierarchical levels—specifically, individual-level absence and colony-level presence—using statistical models. Another aim was to address issues inherent in UAS-based censuses, as they are not merely an extension of the classical field census. For instance, although their low labor intensity enables frequent censuses at short time intervals, the resulting data often exhibit temporal autocorrelation. In addition, object-detection framework needs to be integrated into statistical framework to estimate population size. Using the proposed comprehensive framework, we applied our approach to a breeding colony of black-tailed gulls (*Larus crassirostris*).

Through this application, we estimated population size and evaluated limiting factors, required sample sizes, and the trade-off in sampling effort between UAS-based aerial census and the individual-level tracking. These evaluations allowed to us to assess the applicability of this method across different populations.

## Methods

### Assumptions for applying the UAS and biologging integration framework UAS-based count data

UAS-based aerial censuses are conducted multiple times and yielding count data on the number of individuals detected at time *t*, *A_obs,t_*.

### Bio-logging-based tracking data

Concurrent with UAS-based aerial censuses, bio-logging-based individual tracking is conducted on multiple individuals. Sampling intervals do not need to be high resolution. Both aerial censuses and individual tracking must be repeated at same multiple time points.

### Population

We assumed that the population was closed, with neither immigration nor emigration of individuals from other populations expected during the study period.

### State-space model to infer the population size *N*

We employed state-space models that explicitly account for temporal autocorrelation in the latent state as well as observational errors (Clark & Bjørnstad, 2004; Kéry & Schaub, 2011). Under the assumption of a closed population, we expect a correlation between *Aobs* and *p_t_*, as the following equation holds:

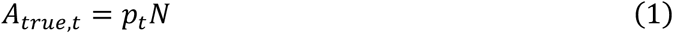

where *A_true, t_* indicates the true number of individuals within the colony. Considering the observational errors, *A_obs,t_* can be expressed using *A_true, t_* as follows:

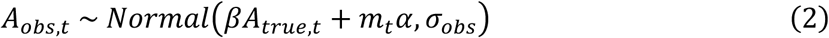

where *p_t_* denotes the probability of an individual being within the colony, and *σ_obs_* represents an observational error. *α* and *β* correct the error of object detection (see Appendix S1: Section S1 for the estimation of *α* and *β*). *m_t_* represents the number of images consisting of an orthomosaic images in a session at time *t*.

Individual-based presence inferred from bio-logging data can be used to estimate *p_t_*. We define the number of individuals tracked at time *t* as *B_t._* We can estimate the number of individuals being within the aerial censused area at time *t* as *B_in,t_*. The relationship between *B_in,t_* and *B_t_* was expressed in a binomial distribution. Furthermore, *p_t_* and *p_t-1_* have similar values under sampling interval is frequent enough. *p_t_* can be expressed using a smoothing trend model as follows:

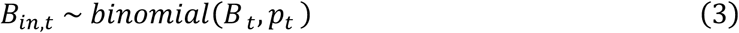

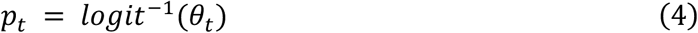

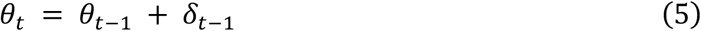

where *θ_t_* is the latent logit-transformed probability of an individual being within the colony at time *t*, i.e., *θ_t_* = logit(*p_t_*). δt-1 represents the difference between *θ_t_* and *θ_t-1_*. By modeling *θ_t_*, we can incorporate temporal smoothness through a state-space formulation while ensuring that *p_t_* remains within the valid probability range of (0, 1).

By estimating all parameters jointly within a Bayesian unified model, we obtain the posterior distribution of *N*. In this paper, we call this model state-space model for *N* (SSMN). For a comprehensive description of the modeling procedure, together with a full list of model parameters and symbols, see Table 2 and Appendix S1: Section S1.

### Case study Data collection

We applied the proposed framework to a breeding colony of black-tailed gulls (*Larus crassirostris*) inhabiting Kabushima Island, Japan (40°32′18″ N, 141°33′27″ E). These birds generally lay eggs in early May, hatching at the end of May and fledging in July. Most colonies consisted of bare ground or low vegetation, except for the central area, which contained shrines, artificial structures, and tall trees (Appendix S1: Figure S1). The birds are central-place forager and *p_t_* was defined as a temporally varying state reflecting temporary emigration (i.e., foraging trip) and subsequent re-immigration.

We conducted 15 flights each during the incubation period in May and chick-rearing period in June at altitudes of approximately 139 m above ground. The flights were automated. Orthomosaic images of the colony were created for each flights, yielding in a total of 30 images (Appendix S1: Figure S1).

We tracked 32 birds with GPS data loggers. The sampling interval was set to 5 min, and data were collected remotely without recapturing. For details on the data collection procedures, see Appendix S1: Section S2 – S5.

### Individual detection in the orthomosaic images using object detection model

We annotated a total of 23,115 adult birds―22,568 grounded birds and 547 flying birds―from 4,209 images with a 640 x 640 pixels size. We divided the annotated dataset into training, validation, and test datasets in ratios of 70, 20, and 10%, respectively. The training dataset included 15,573 individuals from both the grounded and flying birds. Data augmentation with respect to brightness and blurring was conducted. The *n*-model for YOLO-V8 was used (Jocher et al., 2023). We set up 300 epochs and 8 batches in the training, and when the loss function did not decrease in 50 consecutive epochs, training was stopped. We used the default values for the other parameters.

The trained model demonstrated a mean average precision across the two classes was 89.8% (Figure 2A-C; Appendix S1: Appendix Table 1 for other performance metrics). The root mean square error between the number of predicted individuals in the test data and the ground truth revealed was 2.09 per image. Linear regression, with the number of predicted individuals as the dependent variable and the number of annotated individuals as the independent variable, estimated a regression coefficient of 1.07, with a standard deviation (SD) of residuals estimated at 1.18 (Figure 3A). We confirmed that the numbers of images and individuals were sufficient to improve the model (Appendix S1: Figure S2).

**Figure 2.**
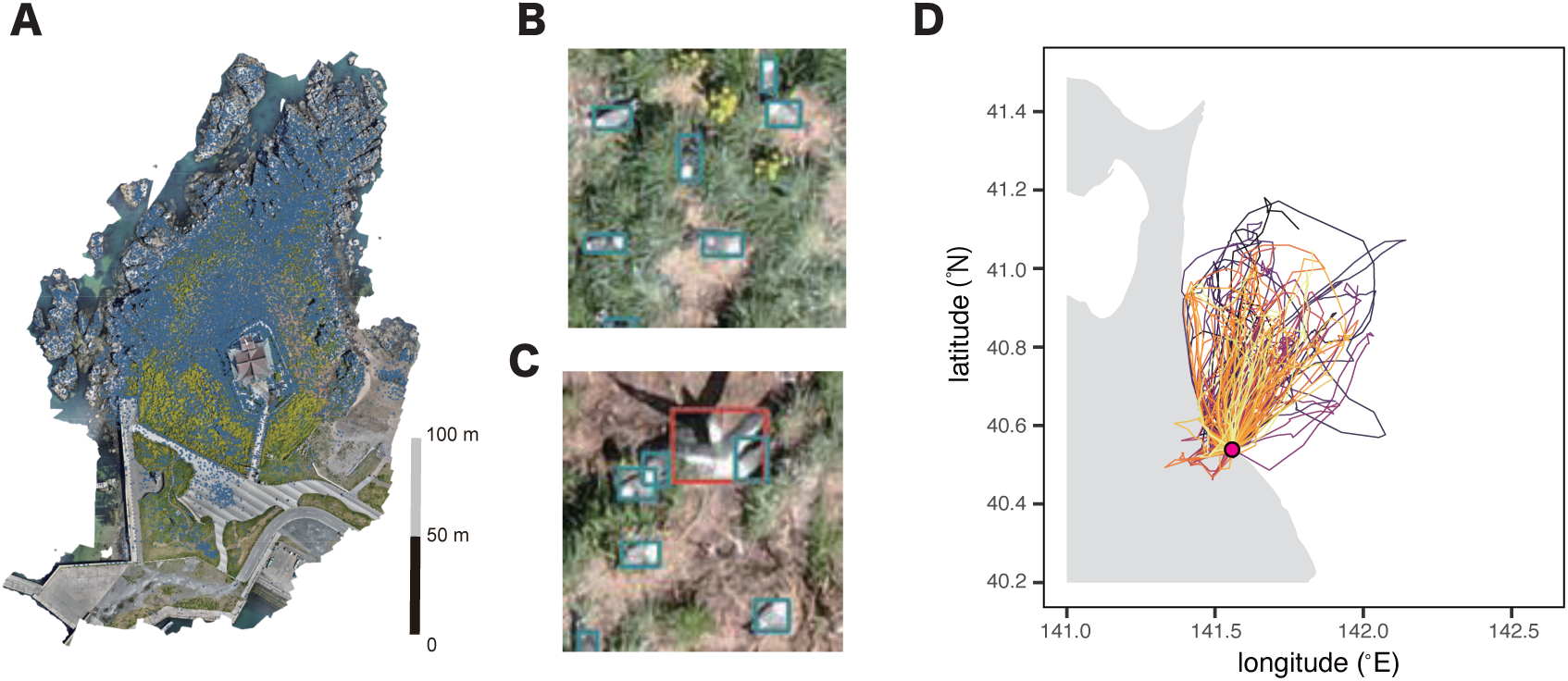
Example of UAS and bio-logging data. **A**: The distribution of detected individuals in the orthomosaic image (blue dots). **B, C**: Partial detection results applied to the orthomosaic image, with blue bounding boxes indicating grounded birds and red bounding boxes indicating flying birds. **D**: Example of bio-logging data derived from one individual for 56 days. Individuals embark on foraging trips mainly to the ocean and return to the colony. The red dot represents the field site.

**Figure 3.**
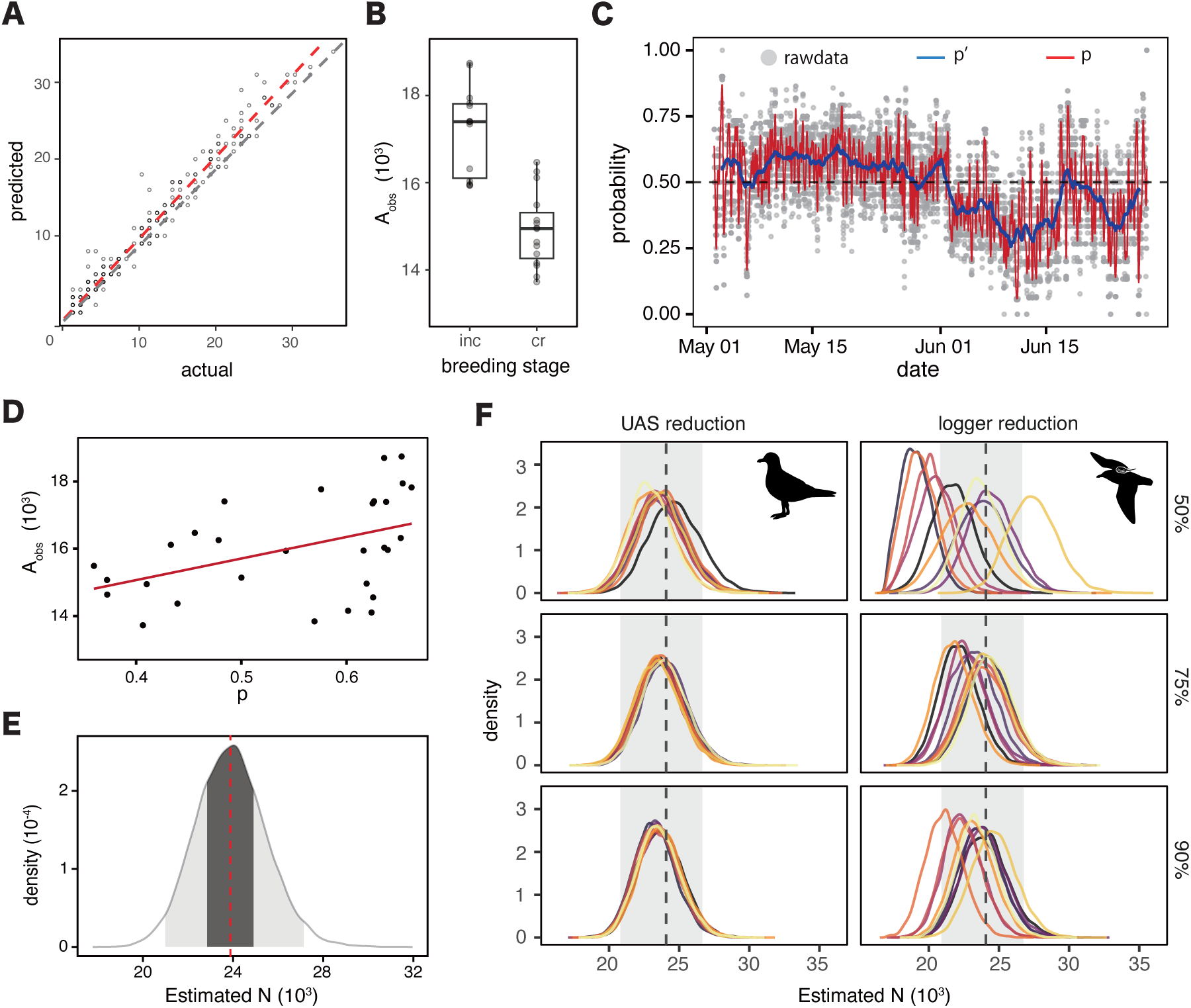
Statistical results of object detection, bio-logging, and state-space models. **A**: The relationship between actual and estimated values in 640 × 640 pixels sized images used for annotation and training. The red dashed line represents the *y = x* line, where points on this line indicate no difference between the estimated and actual number of individuals in each image. The red dashed line represents linear regression results with actual individual counts as the independent variable, suggesting that the object detection model tends toward slight overestimation due to the positive coefficient. **B**: Seasonal variation of number of detected individuals (*Aobs*) between the breeding stages. *Aobs* decreased during the chick-rearing period (cr) compared with in the incubation period (inc). **C**: Temporal variation in *p* and *p′*, which indicates the probability of an individual being in the colony and the proportion of individuals in the colony respectively. The black dashed line indicates a value of 0.5. The red line indicates the estimated *p* via SSMp at each time point. The gray dots indicate the observed values of *p′* at each time point. The blue line represents the moving average of *p′*. The trend of *p′* is around 0.6 but shows a slight decline entering June. **D:** Relationship between *Aobs* estimated by the aerial census and *p* estimated by SSMp. A significant correlation was observed between *Aobs* and *pt* (*p* = 0.03). The line represents the linear regression line. **E**: Distribution of the population size *N* estimated by SSMn. The dashed line indicates the median at 23887, the light grey area represents the 95% credible interval (CI) ranging from 20971 to 27142, and the dark grey area represents the 50% CI from 22838 to 24933. **F**: Posterior distribution of *N* in SMMN using sub-sampled datasets. Each distribution represents the posterior distribution obtained from a randomly sampled dataset, with different colors indicating estimates derived from different samples. The black dasched line represents *N* estimated in the full dataset, and the grey area indicates the 95% CI in the full dataset.

**Table 1.**
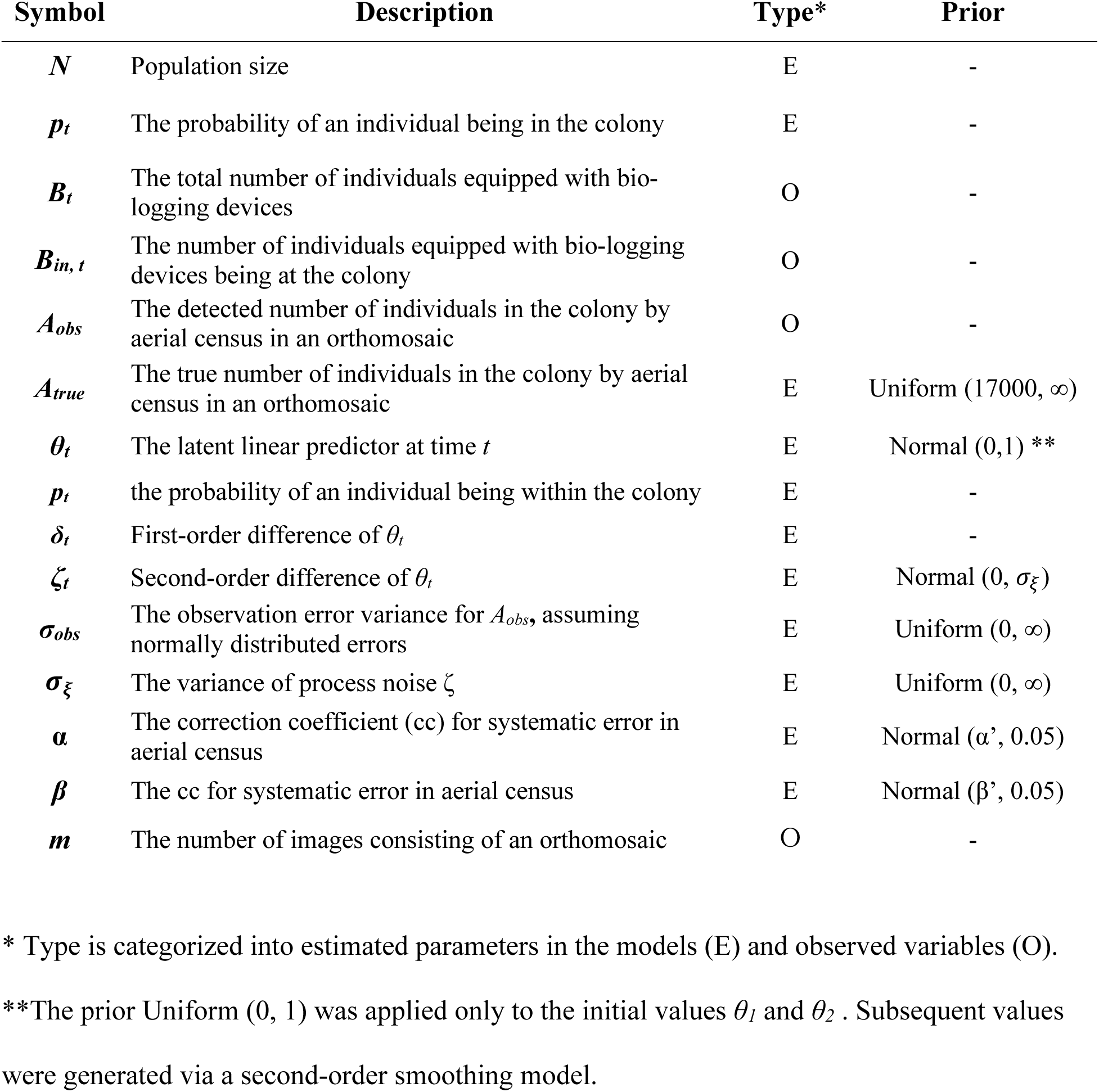
Summary of symbols, definitions, and prior distributions used in the state-space model.

We applied slicing-aided hyper inference (SAHI) to perform predictions on the orthomosaic images (Akyon et al., 2022). Reportedly, SAHI is effective when the target object crosses the edge of images and when the sizes of the images for training and prediction largely differ. Prediction with SAHI revealed 16,119 ± 1,546 (mean ± SD) individuals in one session and that without SAHI indicated 16,454 ± 1,599 individuals, indicating that SAHI suppresses overestimation possibly caused by the double counting of individuals crossing image boundaries (Appendix S1: Figure S2).

In the comparison between May and June, corresponding to the incubation and chick-rearing periods, respectively, the total number of detected individuals (*A_obs, t_*) was lower during the chick-rearing period in June (incubation: 17,316 ± 1,068; chick-rearing: 14,922 ± 861, *p* < 0.0001, Mann-Whitney U test) (Figure 3B). During the chick-rearing period, the number of flying birds was higher (*p* = 0.004, Mann-Whitney U test), whereas the number of grounded birds was lower (*p* < 0.001, Mann-Whitney U test) (Appendix S1: Figure S5).

### Estimating proportion of present individuals in the colony

We resampled GPS locations at 10-min intervals. Each data point was categorized as either ‘in’ or ‘out’ of a 180-m horizontal distance from the central point of the colony, which corresponds to the census area covered by the UAS. We defined *p_t_′* as the simple proportion of tagged individuals that were classified as ‘in’ the colony at time *t*. The value of *p_t_’* was approximately 0.6 during the incubation period, and subsequently decreased to the chick-rearing period (Figure 3C). This pattern aligns with well-documented behavior of gull species, where either parent incubates the eggs and the other leaves the colony for a foraging trip during the incubation period (Beer, 1965).

### Correlation between the probability of an individual being in the colony and the number of individuals detected in the aerial census

Before estimating the posterior distribution of *N*, we evaluated whether the assumptions of our framework were satisfied by constructing a state-space model for *p_t_* (SSMp), where *p_t_* was estimated independently of the aerial census data (Appendix S1: Section S1). We set 20,000 iterations with 4 chains and confirmed that 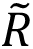 was below 1.1 for the estimated parameters. Estimated *p*_t_ was correlated with the number of individuals in the orthomosaic image *A_obs, t_* (*r* = 0.40, *p* = 0.03, Pearson’s correlation test) (Figure 3D). Although not conclusive, this correlation is consistent with the assumptions of our conceptual model and observation design.

### Estimating population size

Following the SSMp, we estimated *N* using SSMN with 20,000 iterations and four chains and confirmed that B^C^ was below 1.1 for the estimated parameters. The median of *N* was estimated as 23887 (95% credible interval [CI]: 20971 – 27142; Figure 3E). We also estimated the parameters related to object detection and SAHI. The parameter *α* was estimated as 0.32, whereas *α’* was estimated as 0.30 (Appendix S1: Table S3). There was a small difference between *β* (1.06) and *β’* (1.07) (Appendix S1: Table S3). In addition, the value of *β* was similar to the parameter estimated in the linear regression in the object detection (Figure 3A). This indicates that the effect of objective detection will be consistent through small size images prediction and SAHI. For other estimated values, see Table S1.

### Sample size reduction causes parameter estimation instability

To verify whether our sample size was adequate and to examine the variability in estimations, we generated sub-sampled datasets (50%, 75%, and 90%) and repeatedly estimated *N* with 50,000 iterations and four chains. We confirmed that 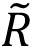 was below 1.1 for the estimated parameters. This sub-sampling analysis revealed that reducing the sample size increased instability in the estimations, with differing levels of impact observed between the reduction of UAS sessions and bio-logging tracks (Figure 3F). When the number of UAS sessions was reduced to 50%, 75%, and 90% of the original dataset, the 95% CIs of all estimations included the estimated *N* for the full dataset (*N_full_*) (Figure 3F). In other words, the posterior distributions of *N* in the sub-sampled datasets did not differ from those in the full dataset. In contrast, in six datasets with a 50% reduction in the number of loggers, the 95% CIs did not include *N_full_* (Figure 3F), although all of the sub-sampled datasets with 90% and 75% reduction included *N_full_* in their 95% CI. To quantify the instability of the estimates under sub-sampling, we calculated the Wasserstein distance between the posterior distribution of *N* obtained from the original dataset and that from each sub-sampled dataset. As a result, reducing the sample size increased the Wasserstein distance from the posterior distribution of the full dataset. Notably, reducing the number of tracked individuals by 90% had a greater impact on estimation uncertainty than reducing the number of UAS sessions by 50% (90% tracks: 746.7 ± 744.4; 50%: UAS 413.3 ± 303.4; Appendix S1: Figure S6).

## Discussion

Since population size serves as an important ecological indicator, it is essential to establish sustainable and accurate population estimation (Callaghan et al., 2024). This study proposes a novel framework for integration of UAS-based count data and bio-logging tracking data. In the framework, aerial censuses and deep learning-based object detection efficiently captured a large number of individuals, revealing fluctuations in their numbers at the colony level and behavioral changes associated with the breeding stage. In parallel with the aerial censuses, bio-logging enabled inferring individual-level presence probabilities within a colony. To bridge these qualitatively and quantitatively different data, state-space models were used to estimate the population size, which could not be achieved using data from only one source. Sub-sampling simulations demonstrated that our sample size provided sufficient accuracy.

Additionally, the simulation results suggested that the number of UAS sessions could be reduced by approximately 50% (15 sessions), although decrease in the number of bio-logged individuals could potentially have a large impact on the estimation results. Thus, these results suggest that allocating sampling effort to bio-logging surveys is more efficient.

### Validity of our framework

Our conceptual model was well supported by the data and the model accuracy. The correlation between the parameters *A_obs, t_* and *p_t_,* which indicate the number of individuals in the colony and the probability of an individual in the colony, respectively, derived independently from UAS and bio-logging techniques, supports the plausibility of our estimated values. This suggests that each data reflect different aspects of the same underlying process.

The integrated model estimated the population size to be approximately 23,000 individuals, which is slightly lower than the results of ground-based nest counts from previous studies using the line transect method between 2007 and 2019 (mean = 15,115 nests, range = 12,042―18,296; Biodiversity Center of Japan, Ministry of the Environment, 2019). In those studies nests were counted within an area of 500 m^2^, corresponding to 0.7% of the total area surveyed in the present study, and these countes were extrapolated to to estimate the population size. Although the results of the present study could suggest a population decline at this site, this apparent difference may instead reflect differences in the methods applied, and the population size may in fact have remained relatively stable. Direct comparisons therefore require caution, given differences in survey season, detection processes, measurement targets, and survey procedures.

### Limitations

Our model relies on the statistical assumption of a closed population. Thus, caution is required when data sampling spans extended periods. To relax this assumption, a robust design sampling scheme can be applied, in which the population is open during the primary sampling periods and closed during the secondary sampling periods (Williams et al., 2002). In our framework, extending the robust design is straightforward. The primary bottleneck is an increased number of aerial censuses, which can be mitigated through automated data collection and processing.

As operational assumptions, the sampled data accurately reflected population characteristics. For instance, when only a subset of individuals uses the area covered by the aerial censuses, the resulting estimates may be biased. Another bias may arise when individuals are present but not detected in aerial images owing to vegetation cover or detector limitations (Couturier et al., 2024). Statistical methods can be employed to adjust for these biases; other solutions may include using high-magnification lenses, infrared cameras, or altering flight patterns (Corcoran et al., 2019). Ensuring that the selection of individuals for bio-logging and that its potential impact is unbiased is also important (Authier et al., 2013; Bodey et al., 2018).

In addition, the colony-level census must adequately capture the target individuals used for individual-level behavioral measurements. Violation of this assumption may increase uncertainty or introduce bias into the estimates. For example, when a target individual is infrequently detected in the colony-level census, the estimated population size *N* may be biased upward. In such cases, incorporating zero-inflated models could provide a potential solution (Lambert, 1992).

A specific limitation of the proposed framework is that handling the number of flying individuals while generating the orthomosaic images can be challenging. Orthomosaic images synthesize still objects that become unstable with excessive movement. In the present study, the overall number of individuals detected decreased in the June session, which included many flying individuals, indicating that the increase in flying behavior could have affected the detection. However, flying individuals accounted for a maximum of 4.5% and a mean of 2.4%, which were relatively low. Considering that *p_t_* also decreased similarly in *A_obs, t_*, this likely did not pose a significant issue. In cases not covered in the current study where the number of flying individuals was high, appropriate measures must be considered. For example, using loggers equipped with accelerometers to understand the precise behavioral state of individuals (Otsuka et al., 2024) could allow for the correction of population estimates by determining the number of individuals in a flying state during UAS photography.

### Perspectives and applications

The data integration using different data streams, but share the identical underlying process, is widely practiced in animal ecology (Benitez-Paez et al., 2021). Previous studies have also suggested that integration of spatial census data and animal movement data not only improve various estimation tasks, but also derive information that cannot be obtained from either one alone (Blackwell & Matthiopoulos, 2024; Thompson et al., 1997; Whitlock et al., 2020; Yamamoto et al., 2015). In line with those studies, the present study highlights the synergistic effects of integrating UAS and bio-logging data using a state-space model that represents both prior knowledge and a conceptual model. In particular, we obtained substantial value of *N* beyond merely combining two different datasets. The accuracy of the data collection process was validated by comparing the parameters obtained from each method. This served to substantiate the validity of the conceptual model of the population, incorporating the expected behaviors of the individuals studied.

Sub-sample simulations highlighted the differences between these data domains. Bio-logging-based tracking data reveals detailed individual-level behavior using a relatively small sample size on a global geographic scale, whereas data collection using UASs, although limited in the spatial range that can be censused, excels in handling the data of multiple individuals using image-based data (Christie et al., 2016). While the UAS-based censuses demonstrated robust results against sample size reduction, the reduction of bio-logging samples decreased model stability, indicating heightened uncertainty and potential overfitting. This is intuitively understandable, as a focus on the detailed behavior of a few individuals may disproportionately influence behavioral interpretations at the individual level. Therefore, in similar situations for the present study, we conclude that when task allocation is required, greater effort should be invested in bio-logging. Understanding the differences between these methodologies can lead to further improvements in efficiency and accuracy and may also broaden their scope of application.

### Novelty of our framework

Absolute population size is particularly important for practical management decision-making. For example, it underpins the assessment of extinction risk, evaluation of environmental carrying capacity, and the setting of quantitative management targets (Callaghan et al., 2024).

While estimating absolute population size typically entails high human effort and strong assumptions (Callaghan et al., 2024), our framework enables the estimation of absolute population size by integrating multiple data sources at relatively low effort.

One novel aspect of our framework is the simplicity of data collection, which may facilitate its application in broader management contexts. UASs conduct a census over a continuous area while detectability was independently estimated using individual-level movement data. Unlike many population estimation approaches (Efford, 2004; Royle, 2004), our framework does not require to dividing the study area into multiple sites or grids. The number and spatial size of such grids can strongly affect estimation results, yet defining optimal grid parameters remains challenging (Sutherland, 2006).

Compared with previous studies in seabird population estimation, our method is advantageous for evaluating the estimation uncertainty. Errors should be considered when estimating breeding nests and individual counting because these surveys must include observation errors and spatiotemporal variation (Frederick et al., 2003). However, the estimation of uncertainty often lacks a solid foundation in general methods because the census may be difficult to conduct multiple times due to the high human effort of a single survey (Wilhelm et al., 2020). Conversely, our study allowed for repeated data sampling with relatively small logistic effort despite the larger census area covering the entire colony, enabling the calculation of annual errors.

An additional advantage of our framework is its capacity for reanalysis as technologies advance. Because census data are derived from single images, future improvements in orthomosaic generation and object-detection models will enable comprehensive reanalysis of the dataset. This flexibility will facilitate deeper insights into population dynamics and enhance adaptive management decision-making.

### Applicability of our framework

The proposed framework has broad applicability to other seabird colony as 95% of seabird species are colonial and typically exhibit surface nesting, except for most species in the Procellariidae family (Schreiber & Burger, 2001; Wittenberger & Hunt, 1985; Young & VanderWerf, 2022). Although our study focused on open environments and highly gregarious, surface-nesting central-place foragers, these traits are not essential, as the key is observing a subset of the population while accounting for individuals inside and outside the census area. For example, this method can be adapted to seabirds that nest on cliffs by modifying UAS flight patterns (Bishop et al., 2022; Brisson-Curadeau et al., 2017). Even for species that nest in rock crevices or burrows, where censusing within their breeding colonies may be challenging, censusing aggregations on the sea surface is feasible. Furthermore, the method can be adapted to other avian species and taxonomic groups, such as colonial marine mammals or other colonial vertebrates, that share similar features.

In rapidly changing environments, exploring sustainable and reliable population estimation methods is essential (Wich & Piel, 2021). The proposed framework is comparatively sustainable with lower observer biases and human effort. Although attaching data loggers requires expertise and skill, the massive deployment of loggers is unnecessary. The autopilot and data analysis capabilities of UASs allow individuals without special skills or training to collect and analyze data through relatively simple protocols. Minimizing observer biases through imaging or direct observations will lead to efficient and sustainable population monitoring. Deep learning often requires advanced programming skills; however, recent developments in codeless programming and generative AI may simplify code operations and maintenance. Although we have emphasized the sustainability of the proposed method, this advantage has the potential to be leveraged to enhance the temporal resolution of monitoring in the future. Realizing this potential will require further methodological and model extensions, but improving temporal resolution will deepen our understanding of temporal population dynamics and ultimately support more informed conservation decision-making.

The proposed integrative approach may facilitate more effective conservation management by reducing human effort and requiring minimal technical expertise for analysis, potentially making it a sustainable solution for population estimation. As multiple data sources become increasingly available, approaches for integrating these diverse data streams are likely to advance over the next decade, enabling new insights into ecosystem processes. Moreover, these methods will become increasingly accessible to practitioners involved in management and monitoring. It is therefore essential that practitioners have a solid understanding of how data are collected and analyzed, as this knowledge is fundamental to recognizing the limitations and potential biases of each study.

## Acknowledgments

We are grateful to Drs. Kenta Ogawa and Ryoma Otsuka for their analytical advice. We thank Taro Okamura for supporting the preparation of Figures and Chihiro Kinoshita for illustrating the seagulls and the UAS in the Figures. Fieldwork was supported by the residents of Hachinohe City.

## Author contributions

SI, YM, YG, KY conceptualized the study. SI, YM, HS conducted the field experiments.

SI, YM, HS, YG analyzed the data. All authors contributed to the writing of this paper.

## Notes

https://github.com/sotainoue/Population-size-estimation-using-UAS-and-bio-logging-data

### Competing Interest Statement

The authors have declared no competing interest.

## References

1. Akyon, F. C., Altinuc, S. O., & Temizel, A. (2022). Slicing Aided Hyper Inference and Fine-tuning for Small Object Detection. In arXiv [cs.CV]. arXiv. http://arxiv.org/abs/2202.06934

2. Anderson, K., & Gaston, K. J. (2013). Lightweight unmanned aerial vehicles will revolutionize spatial ecology. Frontiers in Ecology and the Environment, 11(3), 138–146.

3. Authier, M., Péron, C., Mante, A., & Vidal, P. (2013). Designing observational biologging studies to assess the causal effect of instrumentation. Methods in Ecology and Evolution / British Ecological Society. 10.1111/2041-210X.12075

4. Beer, C. G. (1965). Clutch Size and Incubation Behavior in Black-Billed Gulls (Larus bulleri). The Auk, 82(1), 1–18.

5. Benitez-Paez, F., Brum-Bastos, V. da S., Beggan, C. D., Long, J. A., & Demšar, U. (2021). Fusion of wildlife tracking and satellite geomagnetic data for the study of animal migration. Movement Ecology, 9(1), 31.

6. Biodiversity Center of Japan, Ministry of the Environment. (2019). Monitoring Sites 1000 Seabird Survey Report.

7. Bishop, A. M., Brown, C. L., Christie, K. S., Kettle, A. B., Larsen, G. D., Renner, H. M., & Younkins, L. (2022). Surveying cliff-nesting seabirds with unoccupied aircraft systems in the Gulf of Alaska. Polar Biology, 45(12), 1703–1714.

8. Blackwell, P. G., & Matthiopoulos, J. (2024). Joint inference for telemetry and spatial survey data. *Ecology*, e4457.

9. Boback, S. M., Nafus, M. G., Yackel Adams, A. A., & Reed, R. N. (2020). Use of visual surveys and radiotelemetry reveals sources of detection bias for a cryptic snake at low densities. *Ecosphere (Washington*, D.C), 11(1), e03000.

10. Bodey, T. W., Cleasby, I. R., Bell, F., Parr, N., Schultz, A., Votier, S. C., & Bearhop, S. (2018). A phylogenetically controlled meta-analysis of biologging device effects on birds: Deleterious effects and a call for more standardized reporting of study data. Methods in Ecology and Evolution / British Ecological Society, 9(4), 946–955.

11. Brisson-Curadeau, É., Bird, D., Burke, C., Fifield, D. A., Pace, P., Sherley, R. B., & Elliott, K. H. (2017). Seabird species vary in behavioural response to drone census. Scientific Reports, 7(1), 17884.

12. Callaghan, C. T., Santini, L., Spake, R., & Bowler, D. E. (2024). Population abundance estimates in conservation and biodiversity research. Trends in Ecology & Evolution, 39(6), 515–523.

13. Christie, K. S., Gilbert, S. L., Brown, C. L., Hatfield, M., & Hanson, L. (2016). Unmanned aircraft systems in wildlife research: current and future applications of a transformative technology. Frontiers in Ecology and the Environment, 14(5), 241–251.

14. Clark, J. S., & Bjørnstad, O. N. (2004). Population time series: Process variability, observation errors, missing values, lags, and hidden states. Ecology, 85(11), 3140–3150.

15. Corcoran, E., Denman, S., Hanger, J., Wilson, B., & Hamilton, G. (2019). Automated detection of koalas using low-level aerial surveillance and machine learning. Scientific Reports, 9(1), 3208.

16. Corcoran, E., Winsen, M., Sudholz, A., & Hamilton, G. (2021). Automated detection of wildlife using drones: Synthesis, opportunities and constraints. Methods in Ecology and Evolution / British Ecological Society, 12(6), 1103–1114.

17. Couturier, T., Gaillard, L., Vadier, A., Dautrey, E., Mathey, J., & Besnard, A. (2024). Airborne imagery does not preclude detectability issues in estimating bird colony size. Scientific Reports, 14(1), 3673.

18. Dunn, M. J., Adlard, S., Taylor, A. P., Wood, A. G., Trathan, P. N., & Ratcliffe, N. (2021). Uncrewed aerial vehicle population survey of three sympatrically breeding seabird species at Signy Island, South Orkney Islands. Polar Biology, 44(4), 717–727.

19. Edney, A. J., Danielsen, J., Descamps, S., Jónsson, J. E., Owen, E., Merkel, F., Stefánsson, R. A., Wood, M. J., Jessopp, M. J., & Hart, T. (2024). Using citizen science image analysis to measure seabird phenology. The Ibis, n/a(n/a). 10.1111/ibi.13317

20. Efford, M. (2004). Density estimation in live-trapping studies. Oikos (Copenhagen, Denmark), 106(3), 598–610.

21. Frederick, P. C., Hylton, B., Heath, J. A., & Ruane, M. (2003). Accuracy and variation in estimates of large numbers of birds by individual observers using an aerial survey simulator. Journal of Field Ornithology, 74(3), 281–287.

22. Gallego, D., & Sarasola, J. H. (2021). Using drones to reduce human disturbance while monitoring breeding status of an endangered raptor. Remote Sensing in Ecology and Conservation, 7(3), 550–561.

23. Hayes, M. C., Gray, P. C., Harris, G., Sedgwick, W. C., Crawford, V. D., Chazal, N., Crofts, S., & Johnston, D. W. (2021). Drones and deep learning produce accurate and efficient monitoring of large-scale seabird colonies. Ornithological Applications, 123(3). 10.1093/ornithapp/duab022

24. Hodgson, A., Kelly, N., & Peel, D. (2013). Unmanned aerial vehicles (UAVs) for surveying marine fauna: a dugong case study. PloS One, 8(11), e79556.

25. Hodgson, J.C., Mott, R., Baylis, S.M., Pham, T.T., Wotherspoon, S., Kilpatrick, A.D., Raja Segaran, R., Reid, I., Terauds, A. and Koh, L.P. (2018). Drones count wildlife more accurately and precisely than humans. Methods in Ecology and Evolution, 9(5), 1160–1167.

26. Jocher, G., Chaurasia, A., & Jing, Q. (2023). Ultralytics YOLO (Version 8.0.0). https://github.com/ultralytics/ultralytics

27. Johnston, D. W., Dale, J., Murray, K., Josephson, E., Newton, E., & Wood, S. (2017). Comparing occupied and unoccupied aircraft surveys of wildlife populations: Assessing the gray seal (*Halichoerus grypus*) breeding colony on Muskeget Island, USA. Journal of Unmanned Vehicle Systems, juvs-2017-0012. 10.1139/juvs-2017-0012

28. Kellenberger, B., Marcos, D., Lobry, S., & Tuia, D. (2019). Half a Percent of Labels is Enough: Efficient Animal Detection in UAV Imagery Using Deep CNNs and Active Learning. IEEE Transactions on Geoscience and Remote Sensing: A Publication of the IEEE Geoscience and Remote Sensing Society, 57(12), 9524–9533.

29. Kendall, C. J., Bracebridge, C., Lynch, E. C., Mgumba, M., Monadjem, A., Nicholas, A., & Kane, A. (2023). Value of combining transect counts and telemetry data to determine short-term population trends in a globally threatened species. Conservation Biology: The Journal of the Society for Conservation Biology, 37(6), e14146.

30. Kéry, M., & Royle, J. A. (2020). Applied hierarchical modeling in ecology: Analysis of distribution, abundance and species richness in R and BUGS: Volume 2: Dynamic and advanced models. Academic Press.

31. Kéry, M., & Schaub, M. (2011). Bayesian Population Analysis using WinBUGS: A Hierarchical Perspective. Academic Press.

32. Krause, D. J., Hinke, J. T., Goebel, M. E., & Perryman, W. L. (2021). Drones Minimize Antarctic Predator Responses Relative to Ground Survey Methods: An Appeal for Context in Policy Advice. Frontiers in Marine Science, 8. 10.3389/fmars.2021.648772

33. Lambert, D. (1992). Zero-inflated Poisson regression, with an application to defects in manufacturing. *Technometrics: A Journal of Statistics for the Physical*, Chemical, and Engineering Sciences, 34(1), 1.

34. Lauret, V., Courbin, N., Scher, O., & Besnard, A. (2025). Integrating individual tracking data and spatial surveys to improve estimation of animal spatial distribution. *Ecosphere (Washington*, D.C), 16(5). 10.1002/ecs2.70283

35. Louzao, M., Bécares, J., Rodríguez, B., Hyrenbach, K. D., Ruiz, A., & Arcos, J. M. (2009). Combining vessel-based surveys and tracking data to identify key marine areas for seabirds. Marine Ecology Progress Series, 391, 183–197.

36. Matthiopoulos, J., Wakefield, E., Jeglinski, J. W. E., Furness, R. W., Trinder, M., Tyler, G., Mccluskie, A., Allen, S., Braithwaite, J., & Evans, T. (2022). Integrated modelling of seabird-habitat associations from multi-platform data: A review. The Journal of Applied Ecology, 59(4), 909–920.

37. McCarthy, E. D., Martin, J. M., Boer, M. M., & Welbergen, J. A. (2023). Ground-based counting methods underestimate true numbers of a threatened colonial mammal: an evaluation using drone-based thermal surveys as a reference. *Wildlife Research (East Melbourne, Melbourne*, Vic.), 50(6), 484–493.

38. Otsuka, R., Yoshimura, N., Tanigaki, K., Koyama, S., Mizutani, Y., Yoda, K., & Maekawa, T. (2024). Exploring deep learning techniques for wild animal behaviour classification using animal-borne accelerometers. Methods in Ecology and Evolution / British Ecological Society. 10.1111/2041-210x.14294

39. Reif, J. (2013). Long-Term Trends in Bird Populations: A Review of Patterns and Potential Drivers in North America and Europe. AORN Journal, 48(1), 1–16.

40. Robinson, J. M., Harrison, P. A., Mavoa, S., & Breed, M. F. (2022). Existing and emerging uses of drones in restoration ecology. Methods in Ecology and Evolution / British Ecological Society, 13(9), 1899–1911.

41. Rockwood, L. L. (2015). Introduction to Population Ecology. John Wiley & Sons.

42. Royle, J. A. (2004). N-mixture models for estimating population size from spatially replicated counts. Biometrics, 60(1), 108–115.

43. Rush, G. P., Clarke, L. E., Stone, M., & Wood, M. J. (2018). Can drones count gulls? Minimal disturbance and semiautomated image processing with an unmanned aerial vehicle for colony-nesting seabirds. Ecology and Evolution, 8(24), 12322–12334.

44. Sardà-Palomera, F., Bota, G., Padilla, N., Brotons, L., & Sardà, F. (2017). Unmanned aircraft systems to unravel spatial and temporal factors affecting dynamics of colony formation and nesting success in birds. Journal of Avian Biology, 48(9), 1273–1280.

45. Schreiber, E. A., & Burger, J. (2001). Biology of Marine Birds. CRC Press.

46. Sutherland, W. J. (ed). (2006). Ecological census techniques: a handbook. https://www.researchgate.net/profile/Richard-Shine/publication/223465582_Reptiles/links/00b7d51b139efbba46000000/Reptiles.pdf

47. Sweeney, K. L., Helker, V. T., Perryman, W. L., LeRoi, D. J., Fritz, L. W., Gelatt, T. S., & Angliss, R. P. (2016). Flying beneath the clouds at the edge of the world: using a hexacopter to supplement abundance surveys of Steller sea lions (Eumetopias jubatus) in Alaska. Journal of Unmanned Vehicle Systems, 4(1), 70–81.

48. Thompson, P. M., Tollit, D. J., Wood, D., Corpe, H. M., Hammond, P. S., & Mackay, A. (1997). Estimating harbour seal abundance and status in an estuarine habitat in North-East Scotland. The Journal of Applied Ecology, 34(1), 43.

49. Tuia, D., Kellenberger, B., Beery, S., Costelloe, B. R., Zuffi, S., Risse, B., Mathis, A., Mathis, M. W., van Langevelde, F., Burghardt, T., Kays, R., Klinck, H., Wikelski, M., Couzin, I. D., van Horn, G., Crofoot, M. C., Stewart, C. V., & Berger-Wolf, T. (2022). Perspectives in machine learning for wildlife conservation. Nature Communications, 13(1), 792.

50. van Andel, A. C., Wich, S. A., Boesch, C., Koh, L. P., Robbins, M. M., Kelly, J., & Kuehl, H. S. (2015). Locating chimpanzee nests and identifying fruiting trees with an unmanned aerial vehicle. American Journal of Primatology, 77(10), 1122–1134.

51. Weinstein, B. G., Garner, L., Saccomanno, V. R., Steinkraus, A., Ortega, A., Brush, K., Yenni, G., McKellar, A. E., Converse, R., Lippitt, C. D., Wegmann, A., Holmes, N. D., Edney, A. J., Hart, T., Jessopp, M. J., Clarke, R. H., Marchowski, D., Senyondo, H., Dotson, R.,…Ernest, S. K. M. (2022). A general deep learning model for bird detection in high-resolution airborne imagery. Ecological Applications: A Publication of the Ecological Society of America, 32(8), e2694.

52. Whitlock, S. L., Womble, J. N., & Peterson, J. T. (2020). Modelling pinniped abundance and distribution by combining counts at terrestrial sites and in-water sightings. Ecological Modelling, 420(108965), 108965.

53. Wich, S. A., & Piel, A. K. (2021). Conservation Technology. Oxford University Press.

54. Wilhelm, S. I., Hedd, A., Robertson, G. J., Mailhiot, J., Regular, P. M., Ryan, P. C., & Elliot, R. D. (2020). The world’s largest breeding colony of Leach’s Storm-petrel Hydrobates leucorhous has declined. Bird Conservation International, 30(1), 40–57.

55. Williams, B. K., Nichols, J. D., & Conroy, M. J. (2002). Analysis and Management of Animal Populations. Academic Press.

56. Wittenberger, J. F., & Hunt, G. L. (1985). The adaptive significance of coloniality in birds. Avian Biology, 8, 1–78.

57. Yamamoto, T., Watanuki, Y., Hazen, E. L., Nishizawa, B., Sasaki, H., & Takahashi, A. (2015). Statistical integration of tracking and vessel survey data to incorporate life history differences in habitat models. Ecological Applications: A Publication of the Ecological Society of America, 25(8), 2394–2406.

58. Young, L., & VanderWerf, E. (2022). Conservation of Marine Birds. Academic Press.

