## Appendix for "Integration of UAS-based spatial surveys and bio-logging tracking enhances precision in population size estimation"

**Appendix S1**

### Integration of UAS-based spatial surveys and bio-logging tracking enhance precision in estimating animal colony population sizes

Sota Inoue, Yuichi Mizutani, Hibiki Sugiyama, Yusuke Goto, Ken Yoda

**This PDF file includes:**

Section S1 to S5
Figures S1 to S6
Table S1 to S3

**Section S1: Detailed description of models**

#### State-space model for *p* (SSMp)

The first model, SSMp, was constructed to estimate the posterior distribution of $p_{t,}$ representing. At time $t$, the total number of individuals equipped with bio-logging devices denoted as $B_t$. Those being within the breeding colony were denoted as $B_{in,\,t}$. The relationship between $B_{in,t}$ and $B_t$ was expressed in a binomial distribution as follows:

$$B_{in,t} \sim binomial(B_t, p_t) \quad (1)$$

Assuming that $p_t$ and $p_{t\text{-}1}$ have similar values because the interval is 10 min, $p_t$ can be expressed using a smoothing trend model as follows:

$$p_t = logit^{-1}(\theta_t) \quad (2)$$

$$\theta_t = \theta_{t-1} + \delta_{t-1} \quad (3)$$

Here, $\theta_t$ is the latent logit-transformed probability of an individual being within the colony at time $t$, i.e., $\theta_t$ = logit($p_t$). By modeling $\theta_t$, we can incorporate temporal smoothness through a state-space formulation while ensuring that $p_t$ remains within the valid probability range of (0, 1).

$$\delta_t = \delta_{t-1} + \zeta_{t-1} \quad (4)$$

$$\zeta_t \sim Normal(0, \sigma_\xi) \quad (5)$$

where $\delta_{t-1}$ represents the difference between $\theta_t$ and $\theta_{t-1}$, and $\zeta_{t-1}$ represents the difference in difference. Note that $A_{obs,t}$ is not used in the model; in other words, the UAS-based count data is not used.

Using a Bayesian framework implemented in rstan (Stan Development Team, 2022), we estimated the posterior distribution of the latent state $p_t$ from $B_{in,t}$ and $B_t$, derived from bio-logging data. We ran four MCMC chains for 20,000 iterations each, with a burn-in of 10,000 and a thinning interval of 4, and confirmed convergence by ensuring that all $\tilde{R}$ values were below 1.01 for the estimated parameters.

**State-space model for N (SSMN)**

Following the SSMp, we constructed a state-space model for the population size $N$ (SSMN). Note that $p_t$ was influenced by $A_{obs,\,t}$ during parameter estimation in the SSMN; thus, the estimated $p_t$ could differ between the SSMp and SSMN models.

We defined $A_{true,\,t}$ as the true number of individuals in the colony at time $t$, and $\sigma_{obs}$ as the observational error. Using these parameters, $A_{obs,\,t}$ can be expressed as follows:

$$A_{obs,t} \sim Normal(\beta A_{true,t} + m_t \alpha, \sigma_{obs}) \tag{6}$$

$$\beta \sim nomal(\beta', 0.05) \tag{7}$$

$$\alpha \sim nomal(\alpha', 0.05) \tag{8}$$

$\alpha$ and $\beta$ corrects the error of object detection. $m_t$ represents the number of images consisting of an orthomosaic images in a session at time $t$. We were able to estimate $\alpha'$, $\beta'$ from the test dataset in object detection as follows：

$$A_{pre,i} \sim Normal(\beta' A_{ann,i} + \alpha', \sigma') \quad (9)$$

$A_{pre}$ and $A_{ann}$ represents the number of predicted individuals and the number of annotated individuals in image *i*. Note that image *i* refers a small size (640 x 640 pixels) image, not an orthomosaic. Since SAHI was used to detect individuals in orthomosaics, $\alpha$ and $\beta$ were not identical to $\alpha'$ and $\beta'$, but these parameters were expected to be similar. Therefore, we used $\alpha'$ and $\beta'$ as the mean values of prior distribution of $\alpha$ and $\beta$. We assumed that the population size $N$ remained constant during the survey period. Using $N$ and $p_t$, $A_{true,\,t}$ can be expressed as:

$$A_{true,t} = N \cdot p_t \quad (10)$$

Thus,

$$A_{obs,t} \sim Normal(\beta N p_t + m_t \alpha, \sigma_{obs}) \quad (11)$$

The equations for computing $p_t$ and the procedures for estimating each parameter were identical to those for SSMp.

$A_{obs,t}$ estimated by the UAS-based count data, and $B_{in,t}$ and $B_t$ obtained through bio-logging-based tracking data and those values were used to estimate other values. The prior distribution of $N$ was assumed to be uniform from 17,000 to infinity, where 17,000 represents the minimum estimate based on the UAS-based count data. For a complete list of model parameters and symbols, please refer to Appendix S1: Table S2. To estimate the posterior distribution of $N$, we ran four MCMC chains for 20,000 iterations each, with a burn-in of 10,000 and a thinning interval of 4, and confirmed convergence by ensuring that all $\tilde{R}$ values were below 1.01 for the estimated parameters.

**Section S2: Aerial data collection**

In the aerial census, the altitude above ground fluctuated between approximately 115 and 139 m. An area of 6.4 ha was censused in approximately 25–27 minutes. An autopilot system guided the Mavic 3 drone (DJI, Shenzhen, China), equipped with a 162 mm focal length lens, along a pre-planned rout. Black-tailed gulls usually fly 25 m above the ground [1] and a previous study focusing on the effect of the flight altitude of UASs on bird behavior showed that a flight altitude of 40 m did not affect bird behavior [2]. In the current study, the flight altitudes of the UAS and birds seemed to be largely different, and notable changes in behavior were not detected. We captured approximately 680 images in each session, each with a resolution of 3,840 × 2,160 pixels. The flights were conducted at intervals of either 30 min or 1 h. Sessions were skipped in cases of unsuitable weather conditions, such as rain or strong winds.

**Section S3: Orthomosaic**

Orthomosaic images were constructed from aerial photographs obtained during each session using Metashape (Agisoft LLC, St. Petersburg, Russia), with the Japanese Geodetic Datum 2011 and the Japan Plane Rectangular Coordinate System Zone XVIII (EPSG:6686) used as the reference coordinate system. The areas that could not be successfully merged were excluded from the orthomosaic images. The ground–sample distance was 0.6 cm/pixels. Because of the computational limitations of the computers used, the output from each session was divided into 18 segments, with each segment rendered at 10,000 × 10,000 pixels.

**Section S4: Bio-logging**

GPS data were collected using a PinPoint VHF Solar L (17.6 g; LOTEK, Newmarket, ON, Canada), affixed using a Teflon ribbon harness system [3]. We tracked 32 birds (please refer Appendix S1: Section SX for detailed information). Fifteen of the 32 birds were equipped with the same device in 2022. The number of individuals on which the devices were mounted varied during the study period due to equipment date and device detachment differences. Consequently, data for a minimum of 10 and a maximum of 25 individuals were acquired on the same day (Figure S3). The data acquisition interval was set to 5 min, and data were collected remotely using a VHF commander (LOTEK). We used data from May 2 to June 27, 2024. Considering the potential impact of logger attachment, data from the day of attachment were not used. We resampled GPS locations at 10-min intervals.

Data points are categorized into 'in' or 'out' of a 180-m horizontal distance from the central points of the colony. 180 m corresponds to the census area of the UAS. We defined $p_t'$ as the proportion of tagged individuals that were classified as 'in' the colony at time $t$.

**Section S5: Sub-sampling analysis**

To verify whether our sample size was adequate and to examine the variability in estimations caused by reducing the sample size, we randomly sub-sampled the full dataset and repeatedly estimated it using the SSMN. We constructed six datasets by reducing the number of UAS sessions and bio-logged individuals by 50, 75, and 90%. It should be noted that when reducing the number of UAS sessions, the number of bio-logged individuals did not decrease, and vice versa. Each dataset underwent 10 rounds of estimation using the SSMN. We set 50,000 iterations and four chains. We confirmed that $\tilde{R}$ was less than 1 for the estimated parameters. We compute the Wasserstein distance between the posterior distributions of the parameter $N$ obtained from the original dataset and from sub-sampled datasets.

**Figure S1.**

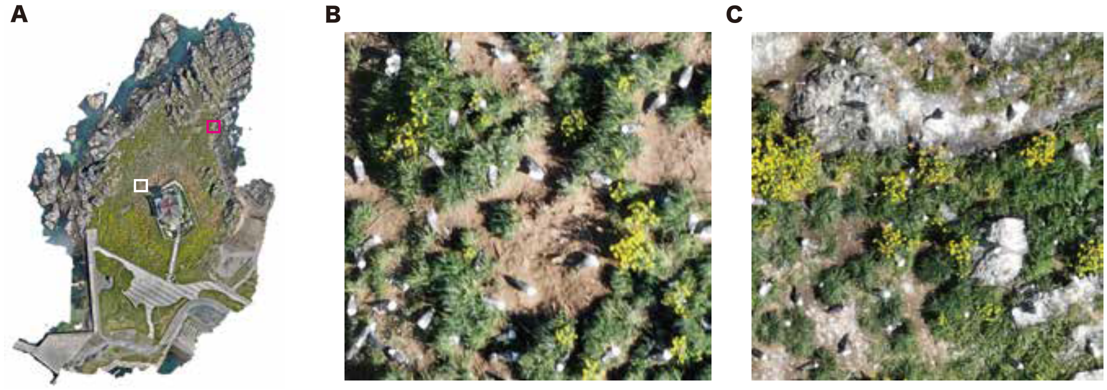


**Figure S1**. Orthomosaic images created by UAS census. A: Example of an entire orthomosaic of the colony. B, C: Images extracted from the orthomosaic image. Black-tailed gulls (*Larus crassirostris*), identifiable by their white heads, are clearly visible in these images.

**Figure S2.**

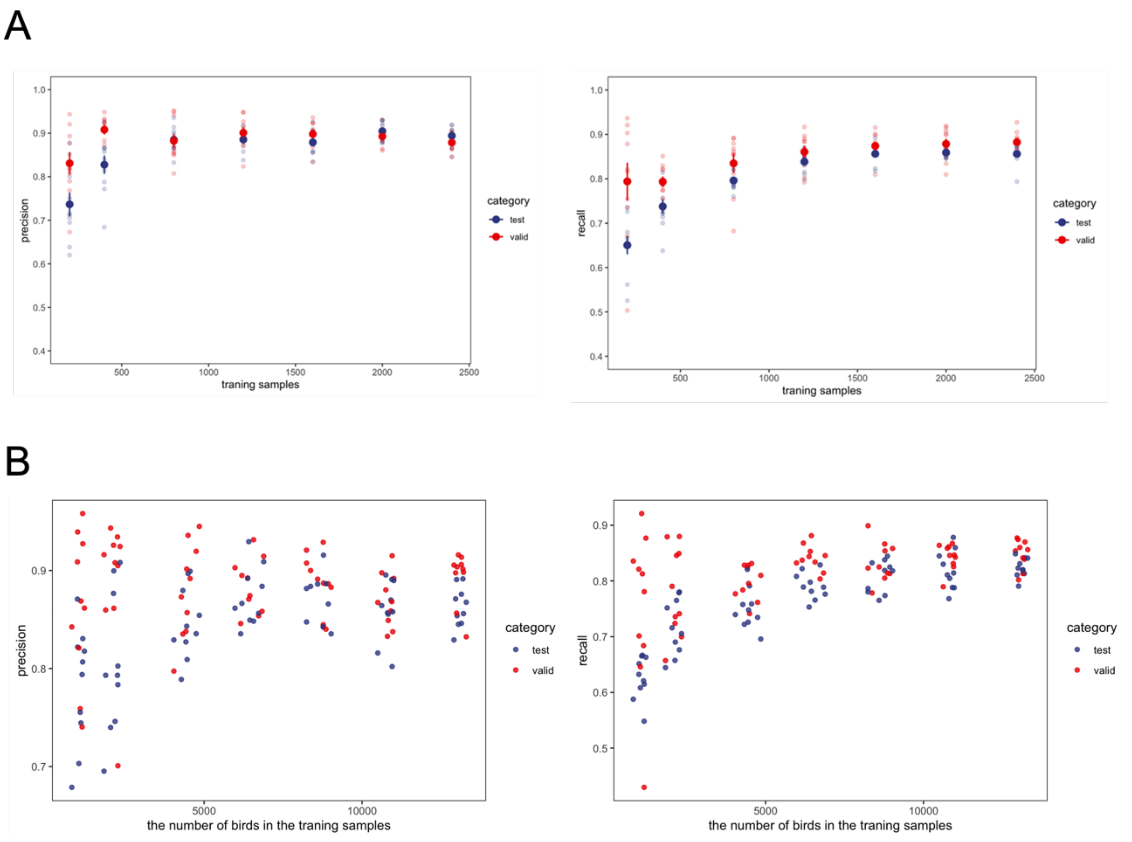


**Figure S2.** To confirm the validity of model-1A, we conducted several experiments. First, we confirmed the validity of the sample size. Different dataset sample sizes, ranging from 200 to 2,400, were created and evaluated. This cycle was repeated 10 times for each dataset. Consequently, the dataset with 2,000 images showed an accuracy similar to the mean average precision of model-1A ($p = 0.74$, Wilcoxon rank-sum test) (Fig. S1A). Next, we examined the influence of variations in the number of individuals per dataset because random sampling of the training data may largely affect the number of individuals in each dataset. The results indicate that the quantity of training data modestly controlled the fluctuation in the number of individuals caused by random sampling. Consequently, increased accuracy was observed with an increased number of individuals within the training dataset. However, the improvement in accuracy plateaued at approximately 10,000 individuals (Fig. S1B). The difference between the maximum and minimum numbers of individuals in the 2,400-image dataset was 389 birds, accounting for approximately 2.4% of the total.

**Figure S3.**

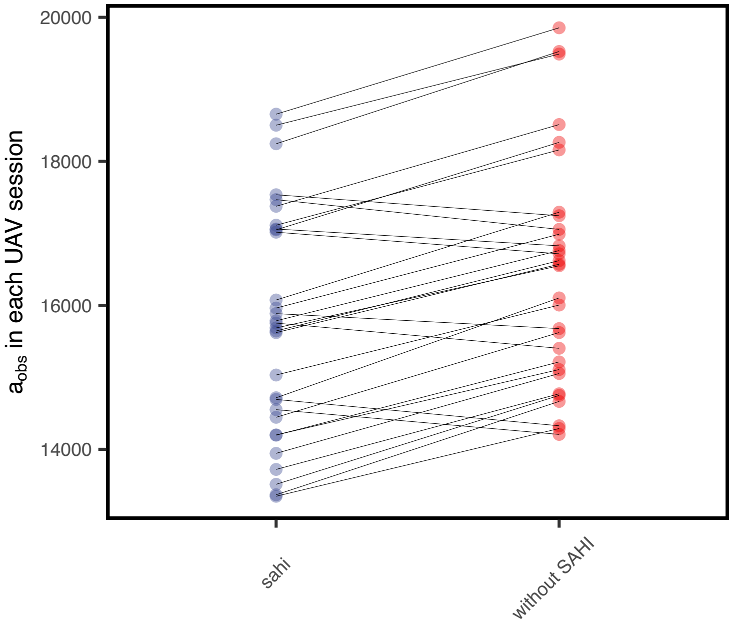


**Figure S3.** $A_{obs}$ in predictions with and without slicing aided hyper inference (SAHI). When SAHI was not applied, $A_{obs}$ was over-estimated.

**Figure S4.**

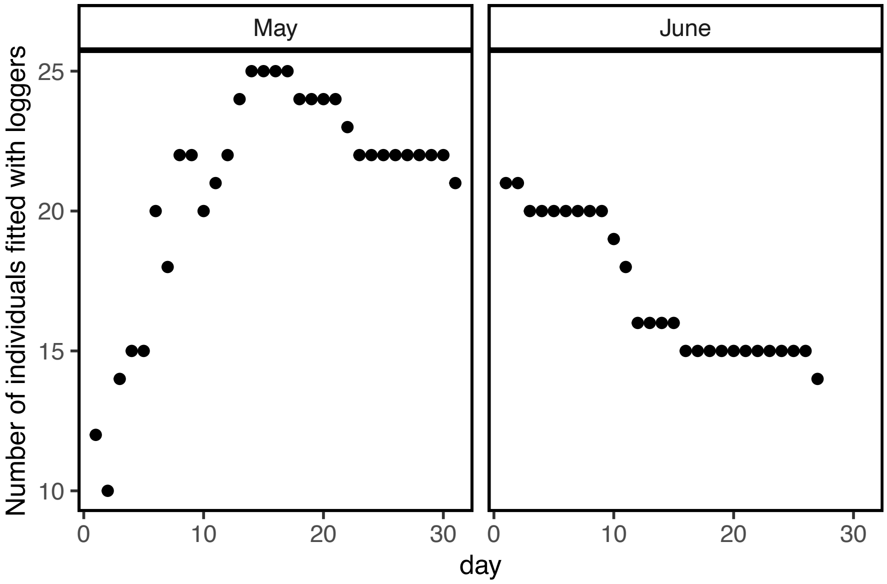


**Figure S4.** Number of individuals fitted with loggers. The number varied during the study period due to differences in equipment date and device detachment.

**Figure S5.**

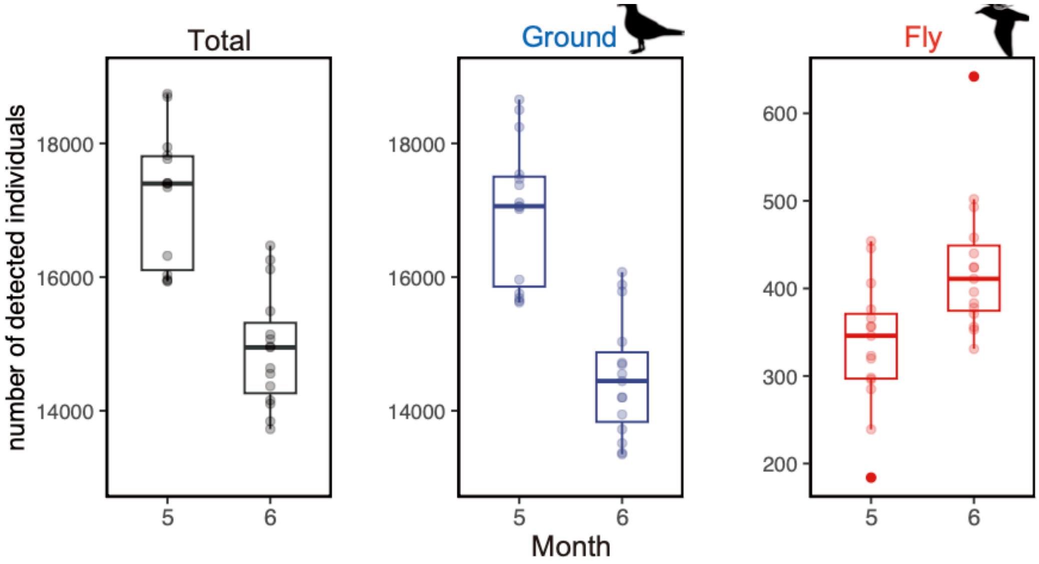


Seasonal variation in each class and detected number of individuals. In the chick-rearing period in June, overall and grounded birds decreased while flying birds increased compared with those in May.

**Figure S6.**

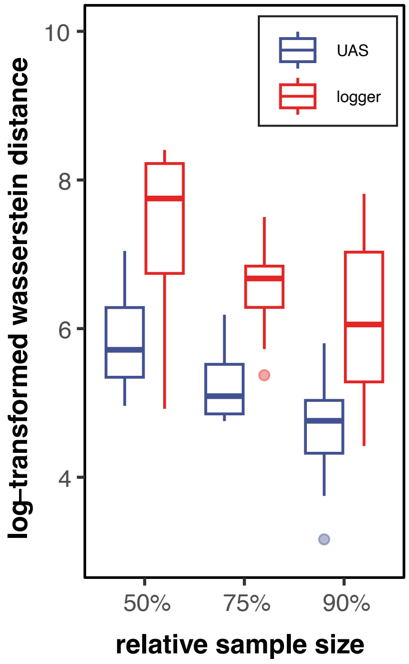


Wassestein distance in sub-sampled datasets. Blue dots indicate the distance for sub-sampled UAS sessions, while red dots represent the distance for sub-sampled logger data.

**Table S1. Evaluations of the object detection model.**

| Images | Class | Instances | Precision | Recall | mAP50 |
|---|---|---|---|---|---|
| | All | 2337 | 0.902 | 0.848 | 0.898 |
| 420 | Grounded | 2287 | 0.858 | 0.877 | 0.908 |
| | Flying | 50 | 0.947 | 0.872 | 0.888 |

**Table S2. Estimated parameters in SSMN.**

**Summary of symbols, definitions, and prior distributions used in the state-space model**

| Symbol | Description | Type* | Prior |
|---|---|---|---|
| $N$ | Population size | E | - |
| $p_t$ | The probability of an individual being in the colony | E | - |
| $B_t$ | The total number of individuals equipped with bio-logging devices | O | - |
| $B_{in, t}$ | The number of individuals equipped with bio-logging devices being at the colony | O | - |
| $A_{obs}$ | The detected number of individuals in the colony by aerial census in an orthomosaic | O | - |
| $A_{true}$ | The true number of individuals in the colony by aerial census in an orthomosaic | E | Uniform (17000, ∞) |
| $A_{pre}$ | The detected number of individuals by object detection model in a small image in the test dataset | O | - |
| $A_{ann}$ | The annotated number of individuals in a small image in the test dataset | O | - |
| $\theta_t$ | The latent linear predictor at time $t$ | E | Uniform (0,1) ** |
| $p_t$ | the probability of an individual being within the colony | E | - |
| $\delta_t$ | First-order difference of $\theta_t$ | E | - |
| $\zeta_t$ | Second-order difference of $\theta_t$ | E | Normal (0, $\sigma_\xi$) |
| $\sigma_{obs}$ | The observation error variance for $A_{obs}$, assuming normally distributed errors | E | Uniform (0, ∞) |
| $\sigma_\xi$ | The variance of process noise ζ | E | Uniform (0, ∞) |
| $\alpha'$ | The correction coefficient (cc) for systematic error in aerial census | E | Uniform (0, ∞) |
| $\beta'$ | The cc for systematic error in aerial census | E | Normal (1, 0.2) |
| $\alpha$ | The cc for systematic error in aerial census | E | Normal (α', 0.05) |
| $\beta$ | The cc for systematic error in aerial census | E | Normal (β', 0.05) |
| $m$ | The number of images consisting of an orthomosaic | O | - |

* Type is categorized into estimated parameters in the models (E) and observed variables (O).

**The prior Uniform (0, 1) was applied only to the initial values $\theta_1$ and $\theta_2$ . Subsequent values were generated via a second-order smoothing model.

**Table S3. Estimated parameters in SSMN.**

| Symbol | Mean | sd | Median | 95%CI | $\widetilde{R}$ |
|---|---|---|---|---|---|
| $N$ | 23917 | 1560.2 | 23887 | 20971 − 27142 | 1.00 |
| $\sigma_{obs}$ | 1857.3 | 380.91 | 1826.1 | 1207.7 − 2670 | 1.00 |
| $\sigma_{\xi}$ | 0.019 | 0.00082 | 0.019 | 0.017 − 0.20 | 1.00 |
| **α** | 0.32 | 0.088 | 0.32 | 0.148 − 0.49 | 1.00 |
| **β** | 1.06 | 0.050 | 1.06 | 0.96 − 1.16 | 1.00 |
| **α’** | 0.30 | 0.071 | 0.30 | 0.155 − 0.44 | 1.00 |
| **β’** | 1.07 | 0.0076 | 1.07 | 1.05 − 1.08 | 1.00 |
| **σ’** | 1.20 | 0.042 | 1.19 | 1.11 − 1.27 | 1.00 |

**References**

1. Park, J.-H., Jeong, I.-Y., Lee, S.-H., Yoo, J.-C. & Lee, W.-S. Changes in Flight Altitude of Black-Tailed Gulls According to Temporal and Environmental Differences. *Animals* **14**, (2024).

2. Rush, G. P., Clarke, L. E., Stone, M. & Wood, M. J. Can drones count gulls? Minimal disturbance and semiautomated image processing with an unmanned aerial vehicle for colony-nesting seabirds. *Ecol. Evol.* **8**, 12322–12334 (2018).

3. Thaxter, C. B. *et al.* A trial of three harness attachment methods and their suitability for long-term use on Lesser Black-backed Gulls and Great Skuas. *Ringing Migr.* **29**, 65–76 (2014).
